# More is better: Improved visual discrimination through local feature diversity in naturalistic objects

**DOI:** 10.64898/2026.01.11.698882

**Authors:** Mina Glukhova, Alisha Crider, Anastasiia Galiagina, Berkutay Mert, Katharine Shapcott, Marvin Weigand, Martha N Havenith, Marieke L Schölvinck

**Affiliations:** Ernst Strüngmann Institute for Neuroscience in cooperation with the Max Planck Society, Zero-Noise Lab, Frankfurt am Main, 60528, Germany

**Keywords:** Visual discrimination, Psychometric curve, Naturalistic stimuli, Morphs

## Abstract

Visual discrimination is traditionally studied with highly controlled, artificial stimuli such as gratings. Yet real-world object discrimination relies on many interacting image dimensions. Here we compared human discrimination performance of oriented gratings with that of natural object morphs, and then asked which stimulus dimensions are most important for morph discrimination. In Experiment 1, participants discriminated between two artificial gratings, two natural grating-like textures, and two natural, morphed stimuli. Morph stimuli were discriminated faster and over a larger stimulus range. In Experiment 2, we separated the contribution of distinct image features, specifically colour, texture, and shape, to the discriminability of the morph stimuli. To this end, we varied two feature dimensions at a time while keeping the third dimension constant. Discrimination was best when colour and texture varied together, resulting in steeper psychometric functions and faster responses than when shape changed. Together, these results suggest a discriminative advantage for natural-object morphs over gratings, driven primarily by local colour and texture information rather than global shape.

## Introduction

In daily life, we navigate through a world full of natural objects that our visual system is exquisitely tuned towards. For example, visual discrimination between the different types of coins in one’s purse allows us to easily pay the right amount at the supermarket, and subtle changes in the appearance of clouds can tell us whether rain is at hand. Yet our knowledge on the discriminative abilities of our visual system is mostly based on artificial stimuli such as gratings (1–3). The reasons for using such highly simplified stimuli are two-fold: they allow for the independent manipulation of various stimulus dimensions (e.g. the orientation, spatial frequency, and colour of a grating), and they enable straightforward quantification of these stimulus dimensions (e.g. grating orientation in degrees).

Using such simplified stimuli, classical studies have found that the visual system is astonishing in its ability to detect very small differences across most stimulus dimensions. In particular, human vision is characterized by extremely fine orientation discrimination, typically less than 1 deg (4, 5). At the same time, it has long been recognised that stimulus dimensions influence each other. For example, the discrimination threshold for the orientation of a grating decreases monotonically with increasing spatial frequency (6). Similarly, shape discrimination depends on blue/yellow or red/green colour vision (7). Therefore, it is likely that the discrimination between natural objects, which differ along many stimulus dimensions simultaneously, results from a (possibly non-linear) interplay between the discriminative abilities of the visual system along different stimulus dimensions.

The dynamics of such natural object discrimination across several stimulus dimensions are difficult to disentangle with lab-designed stimuli like gratings. Instead, research into the visual processing of natural stimuli has so far focused almost exclusively on the processing of natural scenes. For example, photographed scenes with natural spatial frequencies can be discriminated better than scenes with manipulated spatial frequencies (8), and oblique orientations in natural scenes can be discriminated better than horizontal ones (‘the horizontal effect’), which is opposite from what has been found for lab-designed stimuli such as gratings (‘the oblique effect’) (9, 10). Moreover, our sensitivity to luminance contrast is optimized to extract ecologically useful information that is present in the luminance patterns of natural scenes (11), and in spite of their complexity compared to simple lab-designed stimuli, natural scenes activate visual cortex in a very reliable manner (12).

This seminal work has provided the first evidence that the visual system discriminates such stimulus dimensions as spatial frequency and contrast optimally in the range at which they occur in the real world. However, in the real world we typically discriminate between objects, not entire visual scenes. To manipulate the similarity of two objects to each other, morphs can be helpful. Morph images consist of recognizable objects that gradually transform in shape, colour, texture, spatial frequency, etc to that of a different object. Morphs have been used mostly to study shape perception; for instance, it was found that shape discrimination is best around the transition point from one object into another, both in familiar objects (13) as well as in unfamiliar objects (14). Also stimulus attributes, like texture, have been morphed to investigate the perceptual boundaries of various materials (15).

In this study, our aim was two-fold. First, we investigated whether human vision is more precise and accurate in discriminating natural stimuli than classical laboratory stimuli. For this we compared responses to gratings and morphed objects and found that natural objects are indeed processed more effectively by participants. In a second experiment, we then asked to what extent different stimulus dimensions contributed to this improved visual discrimination of morphed natural stimuli. To this end, we independently varied morphed objects along three distinct stimulus dimensions and compared the resulting discrimination performances.

## Experiment 1: Discriminating natural vs artificial stimuli

### Materials and Methods

#### Participants

Experiment 1 included 22 participants (13 female, 9 male, 22 to 41 years of age (mean age 29.27 years)). 8 possessed normal vision and 14 had corrected-to-normal vision (7 with contact lenses, 7 with glasses), 17 were right-handed and 5 were left-handed. Both experiments were approved by the ethics committee from the medical department of the Goethe University in Frankfurt under the study series 2021-252. Participants received written and verbal instructions, completed an eye tracker calibration trial, and completed training trials before continuing to the experimental session.

#### Experimental Setup

Both experiments were conducted in an experimental set-up routinely used for virtual reality (VR) experiments, which consisted of a large spherical dome (diameter 120 cm, 250°; www.fibresports.co.uk) onto which a VR environment was projected via a curved mirror. Participants sat on a pillow with their eyes close to the centre of the dome. A padded, adjustable head and chin rest was used to keep their eyes level and the distance from the dome constant at 73 cm. The VR was created using a custom-made toolbox called DomeVR (16) and consisted of a grassy landscape with mountains in the background and a blue sky overhead, in the middle of which two stimuli (see Experimental Design) were displayed. Responses to the stimuli were given by swiping a GK75-1602B 75mm trackball from NSI (Bilzen-Hoeselt, Belgium) to the right or left, with the right hand.

#### Eye tracking

Throughout each experiment, the eye tracking system iRecHS2 recorded eye movement and pupil dilation at a frame rate of 500 Hz (17). Participants were purposefully not instructed to keep their gaze at the fixation point while the stimuli were displayed. This was done to allow each participant to develop their own viewing strategy.

#### Experimental Design

A trial began with a 500 ms fixation period, followed by a 100 ms delay. Next, two stimuli were shown for 600 ms, between which the participant had to choose within 2400 ms with a leftward or rightward swipe of the trackball. Feedback (correct/incorrect/omission) was given by tones. Omitted trials were repeated at the end of the session to ensure equal numbers of responses across all conditions and difficulty levels.

The stimuli were of three types: circular oriented square-wave gratings (‘artificial gratings’), circles filled with a natural grating-like texture at a certain orientation (‘natural gratings’), or morphs between two types of fish (Fig 1). We chose fish as they are easily recognizable, come in a wide range of sizes, shapes, and colours, and their images can relatively simply be morphed into one another. All stimuli were placed on either a background resembling a grassy field with mountains in the distance (‘mountain background’), or on a homogeneous blue-gray background (‘flat background’). The experiment used a 2AFC procedure, where participants compared one of 12 comparison stimuli against a reference stimulus that was the same on every trial. The correct choice was the grating with the most counterclockwise rotation or the morph stimulus that most resembled the emperor angelfish (Fig 1). Comparing the reference stimulus to itself enabled the analysis of side bias. Each block consisted of 13 trials, presenting all comparison and reference stimuli once in random order. Each block was repeated 10 times, amounting to 780 trials in total (three stimulus types x 13 comparisons x two backgrounds x 10 repetitions). The background at the start of the experiment was individually randomized and then alternated after every three blocks to ensure the participant did not adapt to one background. The order of the blocks was also individually randomized per participant. The position of the reference stimulus (left or right side) was randomized in every trial.

**Fig. 1.**
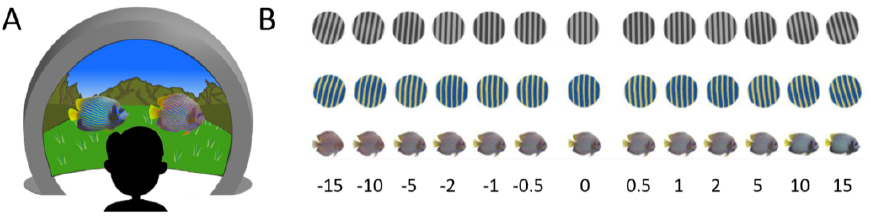
Experimental set-up and stimuli. **A)** Participants were seated inside a large dome, on which a VR and the stimuli were projected. **B)** The stimuli for the three conditions; artificial gratings (top), natural gratings (middle) and morphs (bottom), are shown with the stimulus steps. These steps refer to the orientation differences in the artificial and natural gratings, and to the matched morph steps in terms of Euclidean distances.

#### Stimulus and background parameters

The artificial gratings, natural gratings, and morph stimuli were equated for average luminance, contrast, surface area, and spatial frequency. The morph stimuli contain a multitude of luminances, contrasts, and spatial frequencies, and are therefore difficult to compare to other stimuli. Therefore, we calculated the Euclidean distance between each stimulus pair: the sum of the absolute difference in RGBA values between each corresponding pixel of two images, divided by the number of pixels (see Fig 1B). We calculated the Euclidean distance between each morph pair, and equated the orientation difference between each artificial grating pair and natural grating pair, to match these Euclidean distances (see Suppl Fig S1 for the exact Euclidean distances between all tested stimulus pairs).

In the artificial and natural grating conditions, the reference stimulus had an orientation of 0° (vertical). The 12 comparison stimuli with Euclidean distances matched to the morph stimuli had an orientation of ±0.5°, ±1.0°, ±2.0°, ±5.0°, ± 10.0°, and ±15.0° around the reference stimulus. The morph stimuli ranged from resembling an emperor angelfish to resembling a discus fish, with the reference stimulus looking equally like both fish. For ease of comparison between the conditions, in all figures the 12 morph comparison stimuli are named after the orientation differences (so between ±0.5 and ±15.0), even though these numbers are meaningless for the morphs.

Artificial grating stimuli were created in PsychoPy (psychopy.org) with size 700×700 pixels, spatial frequency of 6, and contrast of 43 percent. The gratings were then uploaded into Krita version 5.0.6 (krita.org), where they were cropped into a circle and their brightness lowered until it matched that of the natural grating stimuli. A random grayscale noise filter was added to the images to increase their Euclidean distance between each other to match that of the morphs. Finally, the image was rotated to create the variations in orientation.

Natural gratings were created using a photograph of an emperor angelfish, the usage rights to which were purchased with a standard usage license (alamy.com). The image was cropped in Krita to a size of 700×700 pixels, a spatial frequency of 6, and rotated to the desired orientations. A desaturation filter was applied to uphold luminance while slightly increasing contrast, and lowering the Euclidean distance between stimuli of different orientations.

The royalty-free images chosen for the morph condition are of an adult emperor angelfish (Pomacanthus imperator)(18) and an adult discus fish (Symphysodon discus) (19). Morphs were generated by first removing the background from the images, and then resizing until the surface area was equal to that of the grating stimuli. With the help of Krita’s desaturation filter, burn tool, and dodge tool, contrast and luminance were slightly adjusted to ensure a match to the other conditions. Next, the G’MIC-Qt plug-in version 3.1.4 (gmic.eu) was used in GIMP version 2.10.32 (gimp.org) to create 400 morph steps between the images. Of those, the morph steps with a Euclidean distance matched to that between the grating stimuli were used in the experiment.

The two backgrounds had a mean luminance of 5.94 cd/m2, recorded using the Konica Minolta CS-100A Chromameter. The RGB value of the flat background was a composite of the average hue and brightness in all stimuli, with RGB values 131, 145, 158 and a luminance of 18.90 cd/m2.

#### Data Analysis

Outcomes of the comparisons (correct/incorrect) were plotted as separate psychometric curves for all conditions and participants. Logistic and Weibull functions were fitted to the curves, and a Bayesian Information Criterion (BIC) was used to evaluate the fits. BIC multiplies the logarithm of the normalized residual sum of squares by sample size. For 20/22 participants, the logistic model best fit the data. Free parameters of this model include y range, y shift, point of inflection, and the maximum slope (Fig 2B). Y range describes the accuracy of responses to the two stimuli, such that consistently providing the correct answer for the extremes of both stimuli would produce a larger y range. Y shift reveals a participant’s response bias, such that a positive y shift would indicate favouring the left side of the stimulus distribution, while a negative y shift arises from favouring the right side of the stimulus distribution. The point of inflection reveals the threshold of discrimination, and the maximum slope of the function indicates performance precision (see Fig 2B).

**Fig. 2.**
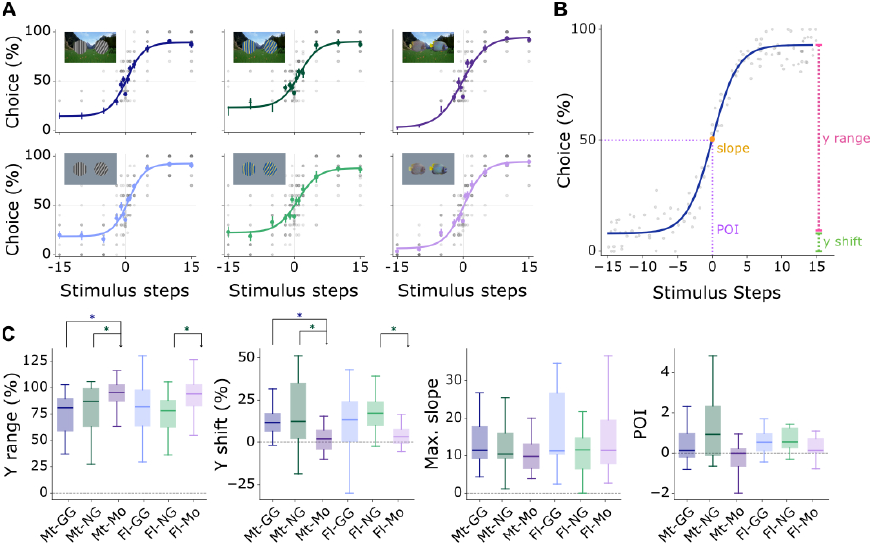
Psychometric functions. **A)** Data and psychometric functions are shown for all conditions. **B)** Parameters describing the psychometric function: y range describes the accuracy of responses at stimulus extremes, y shift reveals response biases, the POI (point of inflection) reflects the discrimination threshold, and the maximum slope indicates performance precision. **C)** Statistical comparison of these parameters for all conditions. Mt, mountain background; Fl, flat background; GG, artificial grating; NG, natural grating; Mo, morph.

The distributions of these parameters were statistically compared across experimental conditions using the Wilcoxon test, FDR corrected for multiple comparisons.

Reaction time (RT) was measured as the time between stimulus presentations and first movement of the trackball, and plotted for each participant, separated by condition and background type. Curve fitting was done on the RT curves using linear, quadratic, and cubic equations. For the curve fitting, the RT to the middle comparison (of the reference stimulus to itself) was left out, as the reaction time here might reflect a very different cognitive process than in the other comparisons. BIC determined that the linear model was the most appropriate representation of the RT data. As such, model parameters for the RT data were slope and intercept. The distributions of curve slopes and intercepts were statistically compared across experimental conditions using the FDR corrected Wilcoxon test.

### Results

#### Psychometric curves

Based on the psychometric curves fitted to participants’ choice probabilities (Fig 2A; see Methods), responses to the morph stimuli differed from the two grating conditions both for flat and mountain backgrounds. These differences were apparent in two curve fit parameters: y range and y shift (Fig 2C). For mountain backgrounds, the y range in morphs is 20.7 percent and 17.6 percent greater than that of natural and regular gratings, respectively (p<0.05, Wilcoxon test, FDR corrected). Similarly, for the flat background, morphs produced a 15.2 percent (p=0.006, Wilcoxon test, FDR corrected) and 11.15 percent (n.s.) greater y range than natural and artificial gratings, respectively. This indicates that morph stimuli were discriminated more accurately than grating stimuli.

Additionally, the morph condition exhibited a significantly smaller y shift than the two grating conditions (for mountain background: p = 0.003 and 0.005, respectively, for flat background: p=0.067 and p=0.006, Wilcoxon tests, FDR corrected). This suggests that response bias was smaller for morph stimuli than gratings. The two grating conditions did not differ along any of these comparisons (Wilcoxon test, p>0.05, FDR corrected). Curve slope and inflection points also did generally not differ across conditions, suggesting that overall discrimination performance was similar (Fig 2C). Thus, while participants were able to discriminate all stimuli with similar precision, morph stimuli produced more accurate and less biased responses than grating stimuli, and the background had no measurable impact on participant performance.

#### Response side bias

Overall, the participants exhibited a negligible left-sided bias of 1.08 percent (range: 17.86 percent left-sided bias to 16.10 percent right-sided bias). This means that the possibility of a response side bias influencing the results can be ruled out.

#### Reaction time

Plotting the reaction times for each stimulus step and condition revealed several interesting aspects. First and foremost, reaction times were generally lower for the morph stimuli than for the two grating conditions (Fig 3A). In all three conditions, there seemed to be a slight effect of task difficulty: stimulus steps close to the reference stimulus led to longer reaction times than stimulus steps far away. In both grating conditions, this effect seemed more pronounced for the positive than for the negative stimulus steps; in other words, the reaction times in the grating conditions were asymmetrical around the reference stimulus (Fig 3A). Lastly, within each condition, reaction times seemed indistinguishable between the mountain and the flat background.

**Fig. 3.**
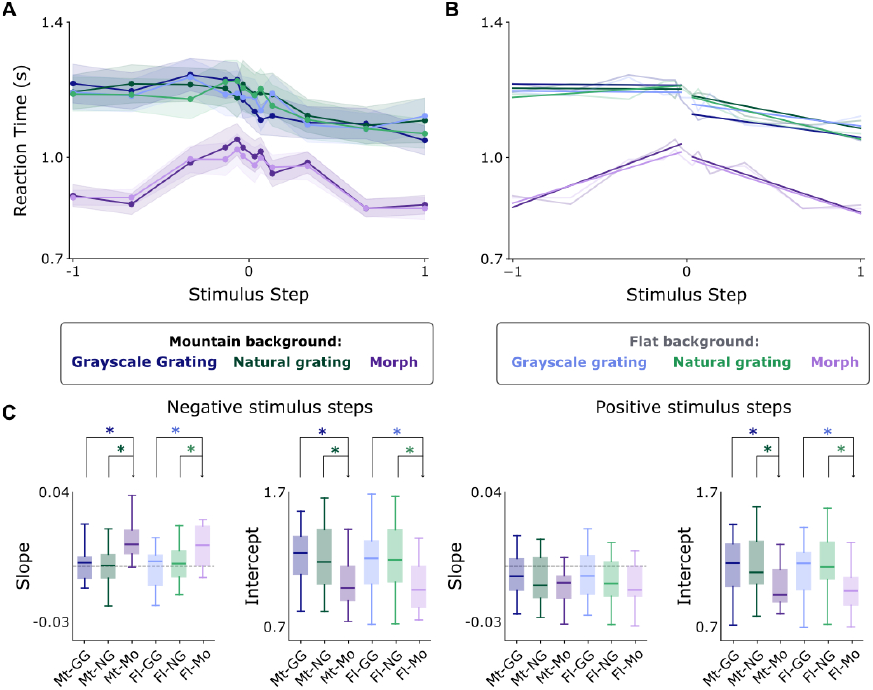
Reaction times. **A)** Reaction times for all three stimulus conditions (see in-figure legend). Lines: Mean reaction times, shaded areas: SEM. **B)** Linear function fitted to reaction times shown in A. **C)** Box plot of the parameters of the linear function, separately for the left (negative) and right (positive) side of the curve. Boxes show median and 25th–75th percentile (IQR); whiskers show 1.5×IQR.

In order to quantify these observations, we fitted linear curves to the two stimulus sides (positive and negative stimulus steps) and each condition and background separately (Fig 3B). The y-intercept of the morph fits was indeed significantly lower than that of the gratings, pointing towards generally lower reaction times (p<0.005 for all conditions, Wilcoxon test, FDR corrected; Fig 3C). Although the slope of all conditions seemed to be non-zero thus pointing to an effect of task difficulty, this was only significant for morph conditions (for all backgrounds and both stimulus sides, p<0.05, Wilcoxon test, FDR corrected). However, task difficulty did impact reaction times to morph stimuli more than to either type of grating in the negative stimulus steps (p<0.05 for both conditions and both backgrounds, Wilcoxon test, FDR corrected). While the differences in slope and intercept between the negative and positive orientations were mostly not significant (apart from the intercept for artificial gratings on a mountain background and the slope for morphs on both backgrounds), thus not statistically confirming the asymmetry around the reference stimulus, it is noticeable that parameters for negative stimulus steps were impacted more significantly by stimulus condition than for positive stimulus steps. Lastly, the average reaction time was not affected significantly by the two background conditions (p>0.05 for all conditions, Wilcoxon test, FDR corrected).

## Experiment 2: Determining relevant stimulus dimension

### Materials and Methods

#### Participants

Experiment 2 included 13 participants (7 female, 6 male, 21 to 51 years of age (mean age 33.41 years)), six of whom had also participated in Experiment 1. Five possessed normal vision and eight had corrected-to-normal vision, ten were right-handed and three were left-handed. For technical reasons, the eyes of the participants could not be tracked in this experiment. Participants received written and verbal instructions and completed training trials before continuing to the experimental session.

#### Experimental Design

To test if different stimulus dimensions contributed with varying extent to the visual discrimination between two morphed natural stimuli, we focused on three basic visual features of natural objects: colour, texture, and shape. As in Experiment 1, we used morphs between an angel fish and a discus fish, and we created 13 stimulus pairs consisting of a reference stimulus and another stimulus, with increasing Euclidean distance between the two stimuli. To facilitate keeping the Euclidean distances equal between the morph pairs, we simultaneously manipulated two stimulus dimensions while keeping the third dimension constant. Thus, we created morph pairs that differed in either 1) colour and texture (keeping the shape constant), 2) colour and shape (keeping the texture constant), or 3) texture and shape (keeping the colour constant) (see Fig 4A). As experiment 1 revealed no differences in psychometric curves and reaction times between flat and mountain backgrounds, we chose to only present the stimuli in Experiment 2 on the mountain background. This allowed us to increase the number of repetitions of each difficulty level within each condition from 10 to 15. The experimental design was otherwise identical to Experiment 1.

**Fig. 4.**
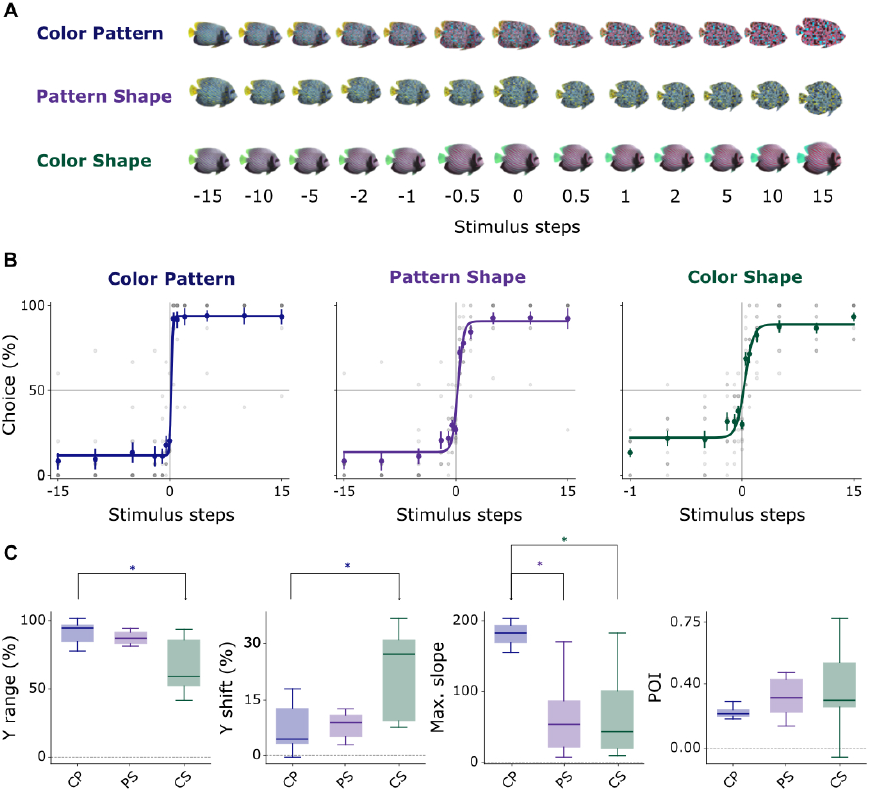
Stimuli and psychometric functions. **A)** Stimuli for the three conditions in Experiment 2. **B)** Psychometric functions for the three conditions. **C)** Statistical comparison of the parameters of the psychometric functions. The labels on x-axis are: CP = Colour x pattern, PS = patten x shape, CS = colour x shape,

#### Stimulus and background parameters

All stimuli were based on the same angel fish and discus fish images as presented in Experiment 1. For morphs along the colour dimension, we used the colour balance tool in KRITA to recolour both fish and then adjusted the saturation and contrast to keep them constant across all stimuli. Colours changed such that yellow tones turned into cyan while blue tones turned into red. For the texture change, we divided the original angel fish into 26×16 sections to produce sections of a reasonable size, and then randomly shuffled and rotated each section (Suppl Fig S2A) to create a different pattern. On the morphs from the angel fish into the discus fish, we used the patterned image as an overlay with increasing opacity (Suppl Fig S2B). For all three conditions, we used the G’MIC-Qt plug-in in GIMP to create 400 morph steps between the images. Of those, the 12 morph steps with equivalent Euclidean distances between the conditions were used in the experiment (Suppl Fig S3).

#### Data Analysis

As in Experiment 1, outcomes of the comparisons (correct/incorrect) were plotted as separate psychometric curves for all conditions and participants. Logistic and Weibull functions were fitted to the curves, and the Bayesian Information Criterion (BIC) was used to evaluate the fits. For all participants, the logistic function best fit the data. The distributions of y range, point of inflection, and the maximum slope of the logistic function were statistically compared using a repeated measurement Wilcoxon signed-rank test, FDR corrected for multiple comparisons. Reaction times were measured, fitted and statistically tested in the same way as in Experiment 1.

### Results

#### Psychometric curves

The slope of the fitted psychometric function can be taken as a measurement for how discriminable the stimulus dimensions are. The largest average slope was found for the ‘colour and pattern’ condition (183), which was significantly larger than the other two conditions (p<0.005, Wilcoxon test, FDR corrected); the slopes of the other two conditions (54 for ‘colour and shape’ and 43 for ‘pattern and shape’) were not significantly different.

The y range (see Fig 2B) was largest for the ‘colour and pattern’ condition (p=0.02 compared to ‘colour and shape’ condition, Wilcoxon test, FDR corrected; the other two comparisons were not significant), implying that this condition was also most accurately discriminated. Y shift was largest for the ‘colour and shape’ condition (p=0.03, Wilcoxon test, FDR corrected compared to ‘colour and pattern’; nonsignificant compared to ‘pattern and shape’), indicating a slight response bias for this condition. There were no significant differences in the point of inflection between the conditions, implying a similar threshold of discrimination for all conditions.

#### Reaction time

In agreement with the psychometric function slope differences, the ‘colour–pattern’ condition yielded the fastest reaction times (lower average RT than both colour–shape and pattern–shape on both sides (p=0.0051), while colour–shape and pattern–shape did not differ (p=0.424), Wilcoxon test, FDR-corrected), implying that the ‘colour-pattern’ condition was not only most accurately, but also fastest discriminated (Fig 5A). In line with Experiment 1, there was no asymmetry in RTs around the reference stimulus for the morphs in Experiment 2. However, the increase in reaction times for the stimuli close to the reference stimulus shown in Experiment 1 was also not present, and overall, all RTs were quite low - in fact, most RTs aligned with the lowest RTs recorded in Experiment 1. Together with the steep psychometric curves, this probably indicates that the task was perceived as quite easy. Linear fits nevertheless showed a stimulus-dependent RT change in most cases (Fig 5B, Wilcoxon signed-rank vs zero slope, FDR-corrected): slopes were significantly different from zero for colour–pattern and pattern–shape on both stimulus sides (p<0.05 for all 4 conditions), and for colour–shape on the positive side (p=0.0037; negative side n.s.) (Fig 5C). There was a slight asymmetry in RTs to the two stimulus sides, with consistently faster responses to the positive than to the negative stimulus steps in all conditions (p<0.05 for all conditions, Wilcoxon test, FDR-corrected). Lastly, there was a small intercept difference for colour–pattern (p=0.0015, Wilcoxon test, FDR-corrected).

**Fig. 5.**
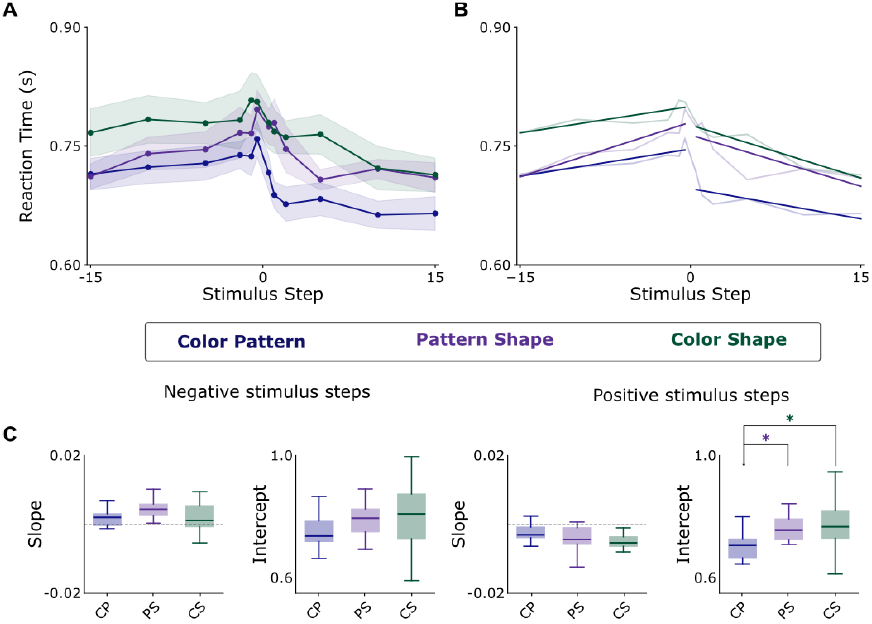
Reaction times. **A)** Reaction times for the three conditions in Experiment 2. **B)** Statistical comparison of the parameters of the linear fit of the reaction times. Box plot of the parameters of the linear function, separately for the left (negative) and right (positive) side of the curve. Boxes show median and 25th–75th percentile (IQR); whiskers show 1.5×IQR.

## Discussion

In this study, we investigated the ability of the visual system to discriminate natural stimuli, by comparing the orientation discrimination of artificial gratings and natural image gratings to the discrimination of morph stimuli. We found that morph stimuli were discriminated better (larger y-range of the psychometric function), and faster (lower RT baseline) than artificial and natural grating stimuli. These effects were independent of the background on which the stimuli were portrayed, despite earlier work showing that the perceived contrast of a luminance patch is suppressed when it is embedded in a real world scene (20). Moreover, this increased performance on the morph stimuli seemed to be due in large part to their colour and texture (steeper slope of the psychometric function and lower reaction times), not their overall shape.

The discriminative advantage of the morph stimuli over the gratings comes despite the vertical orientation of the reference stimulus, thus allowing for the oblique effect: the phenomenon that participants can most quickly and accurately distinguish between gratings when they are at or near cardinal directions (21). No substantial difference in performance was observed between our artificial and natural grating conditions, despite literature suggesting humans discriminate better between gray-scale gratings than blue-yellow gratings (7, 22). The colour combination of the natural grating condition may have decreased subject performance, while the additional visual information given by the natural features (i.e. fish scales or the texture) may have partially rescued this effect.

The lower reactions times for the morphs is in agreement with faster reaction times in natural images compared to synthetic features (23). In all conditions, RTs showed also an effect of task difficulty: discrimination close to the reference stimulus took longer than discrimination far away from the reference stimulus. In the morphs, this effect was most pronounced, which may indicate that humans inherently differentiate more quickly between lifelike creatures than between more abstract objects, like gratings that differ only in orientation. In the two grating conditions, RTs were lower for the left-tilted gratings than the right-tilted gratings. Since the choice for the participant was, ‘which grating tilts more to the left’, it was apparently easier to respond to a grating with a large tilt than to the reference grating with zero tilt (which is the correct answer when it is compared to a right-tilted grating).

Discrimination of the gratings and morphs was directly compared through the use of the Euclidean distance measure. We chose this measure as in our opinion, it provided the most objective comparison between two gratings or between two morphs. However, the Euclidean distance does not take any properties of the visual system, such as its typical responses to contrast or edges, into account; as a result, the actual difference between two objects (gratings or morphs) may be very different from their perceived difference. A model that does take such knowledge of the primary visual cortex into account, is the multiresolution colour model for visual difference prediction (8). This model carries out a multiresolution analysis of two pictures and detects differences in local contrast and colour at each spatial frequency. The model is validated against experimental psychophysics data. Although this model would have better reflected how the visual system discriminates between two natural stimuli, it is optimised for small (near-threshold) differences only and for natural scenes, and would therefore not have provided the most objective way to compare between gratings and morphs.

Why were morph stimuli better discriminated than gratings? An important clue to this comes from the general finding of Experiment 2: when both colour and texture of the fish change (and not the overall shape), all the differences between the various morphs are manifested locally. In other words, when focusing only on a small part of the morphs instead of on the entire morphs, in the ‘colour and pattern’ condition one receives the most information to discriminate between them. The superior discrimination in this condition implies that discrimination is based mostly on local visual information. This is in agreement with the fact that local orientation clues aid discrimination more than global orientation clues, and that local information pooled from various spatial locations yields the best discrimination (24). It is also in agreement with experiments on sensitivity to local natural image regularities: observers can learn to discriminate the statistical regularities of natural image patches from those represented by patch-based probabilistic natural image models after very few exposures, suggesting that the visual system is biased for processing natural images, even at very fine spatial scales (25). In line with this, an increase in spatial frequency eases orientation discrimination (6); at each local ‘spot’, there is simply more information to discriminate between orientations.

These findings suggest that the morph stimuli in the first experiment contained more ‘local’ information to base the discrimination on than the gratings. In our experiments, we equated the average Euclidean distance over the entire stimuli between the conditions, but plotting the Euclidean distance pixel by pixel for the three conditions revealed substantial differences in their distribution: the two grating conditions showed stripes of small and large Euclidean distances, whereas the Euclidean distances in the morph condition were very homogeneous across pixels (Suppl Fig S4). This confirms that indeed, in the morph stimuli, a random local spot contained potentially more information for discriminating between stimuli. Moreover, orientation discrimination might be based on the relation between neighbouring locations (i.e. second order statistics) rather than on local (i.e. first order statistics) measures of differences (26, 27). It might therefore be computed in a fundamentally different way.

Given the differences in higher order statistics between the gratings and the morphs and the corresponding differences in discriminability, it is remarkable that the background had no quantifiable effect on performance. One might hypothesise that the colours in the mountain background may have distracted participants from differences in the stimuli, while the horizon gave orientation cues. In addition, the image statistics of a natural scene are known to influence object recognition, e.g. the presence of animals in a natural scene (28). In contrast, our homogeneous background and our natural scene background gave similar discrimination performance, both for gratings as well as for morphs. This might be partly due to a semantic mismatch between the morphed fish and the mountainous background, which we chose out of practical reasons.

Previous research on natural scenes has shown that it is their natural image statistics that makes them better discriminable. The research presented here adds to this by showing that natural objects are better and faster discriminated than artificial stimuli thanks to their abundance of local difference information. This suggests that the combined processing of the various stimulus dimensions of natural objects boosts the performance of our visual system.

## ACKNOWLEDGEMENTS

We thank Valero Laparra and Jesus Malo Lopez for their helpful input on the Euclidean distance measure.

## Supplementary Figures

**Fig. S1.**
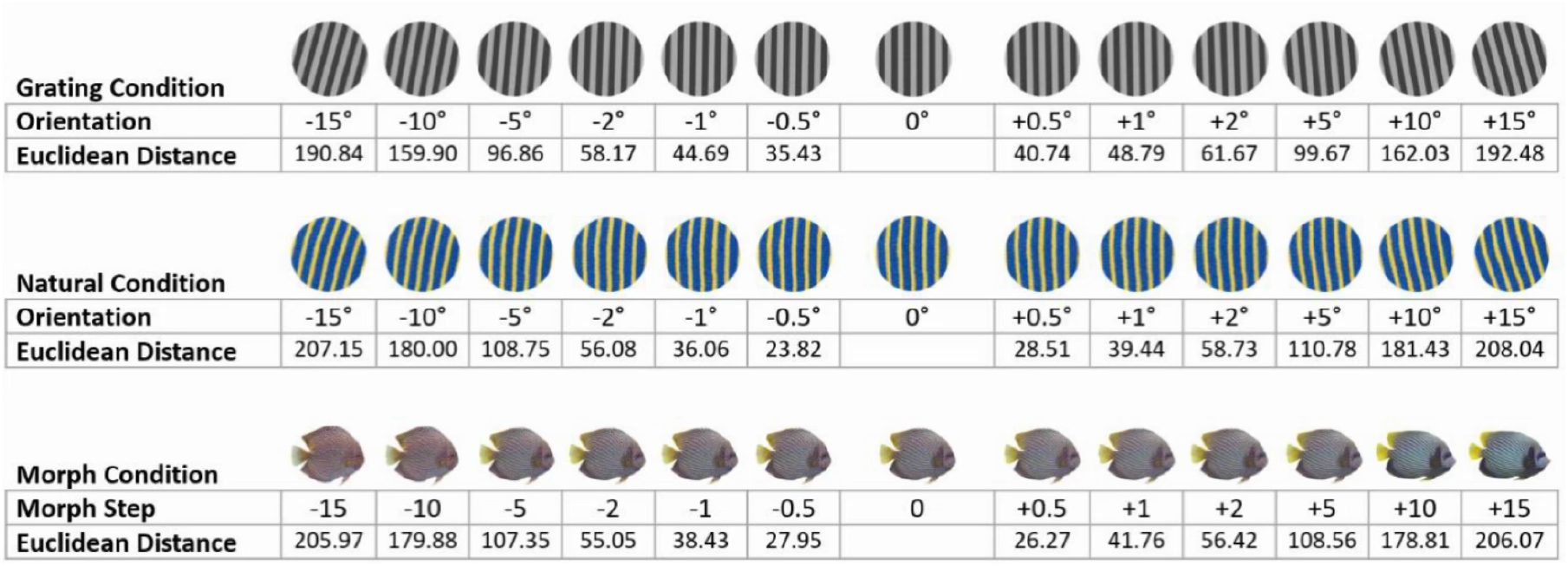
Stimulus distances in Experiment 1. Euclidean distances between each stimulus step and the reference stimulus, for artificial gratings (top row), natural gratings (middle row) and morphs (bottom row).

**Fig. S2.**
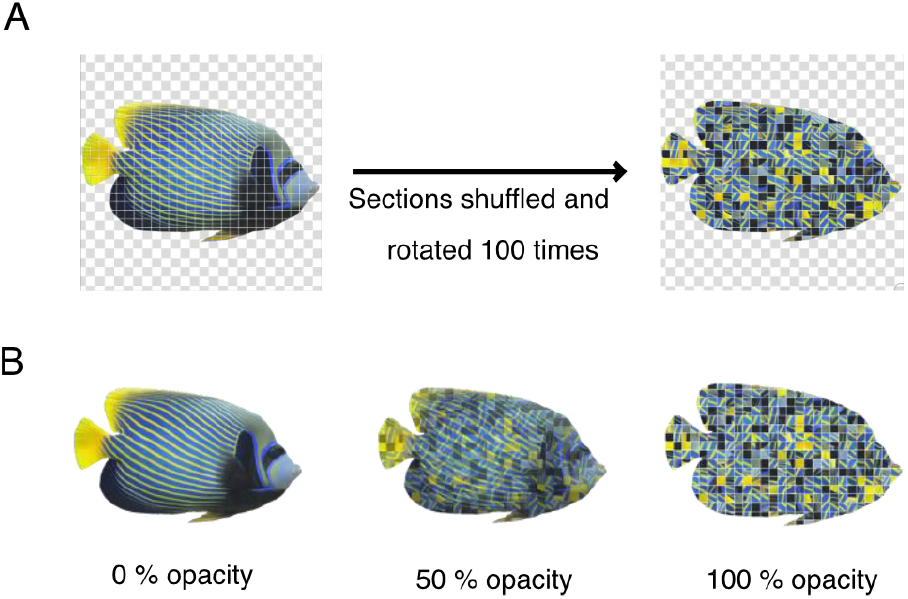
Texture manipulation. A) The original discus fish was divided up into sections, which were shuffled and independently rotated 100 times to create a different pattern on the fish. B) This patterned image was used as an overlay with increasing opacity on top of the angelfish image.

**Fig. S3.**
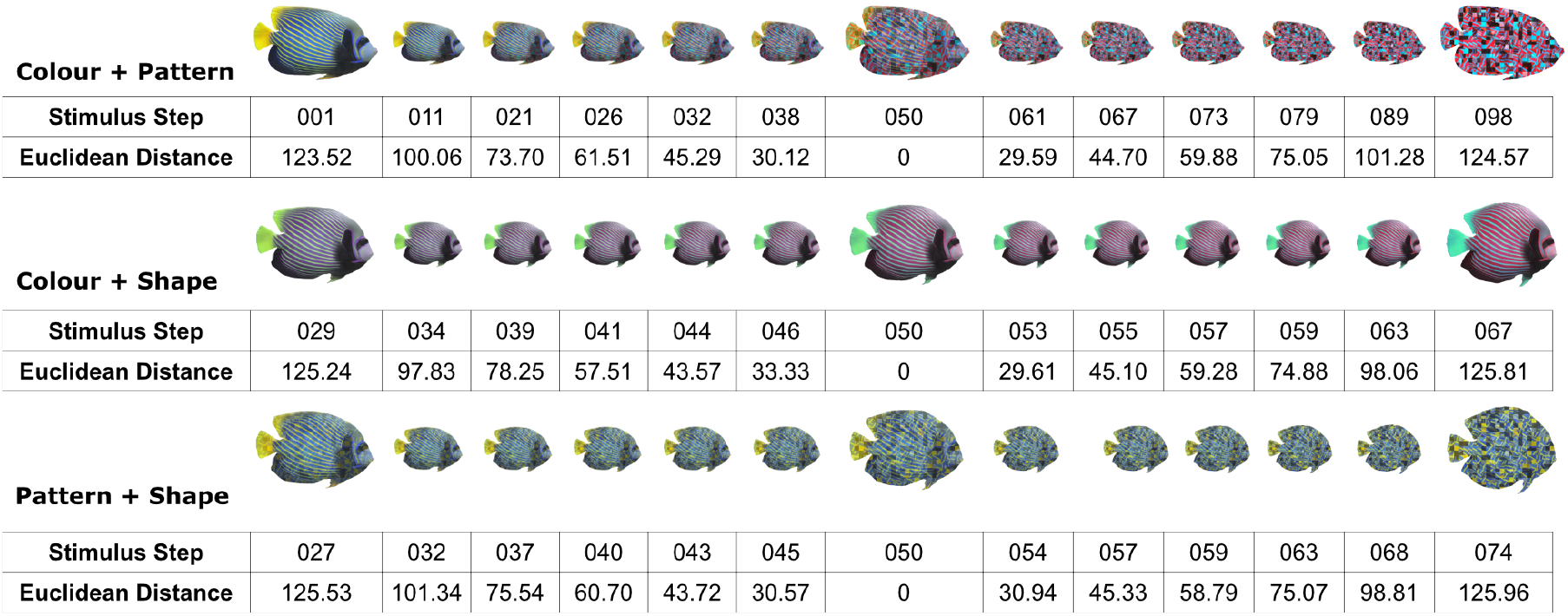
Stimulus distances in Experiment 2. Euclidean distances between each stimulus step and the reference stimulus, for the colour-pattern (top row), colour-shape (middle row) and pattern-shape (bottom row) conditions.

**Fig. S4.**
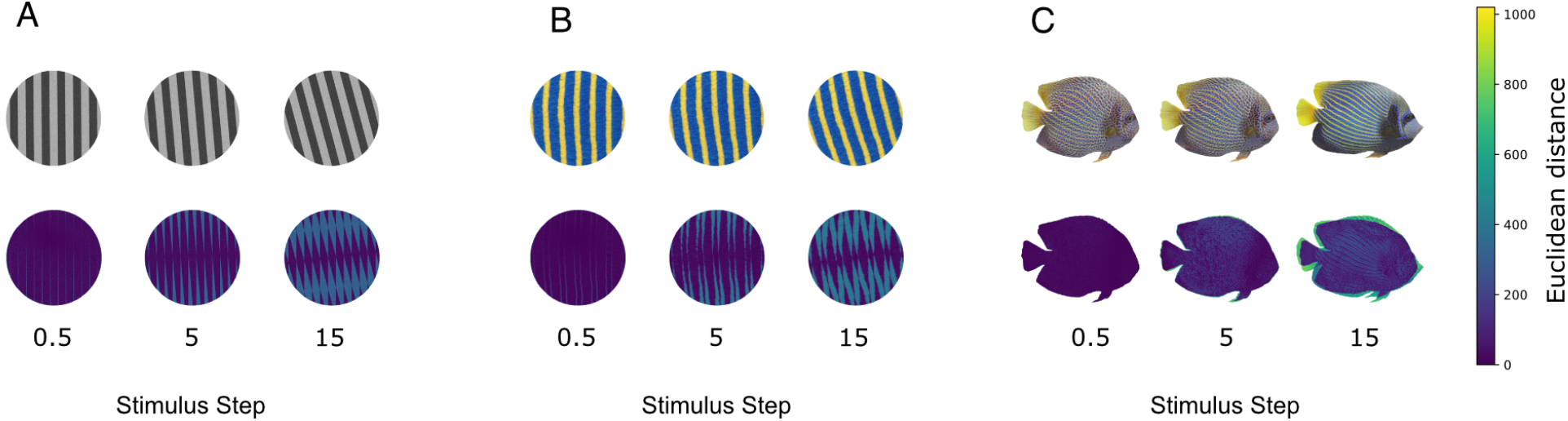
Visualisation of stimulus distances. Pixel-by-pixel Euclidean distances between each stimulus step and the reference stimulus, for: A) the artificial gratings, B) natural gratings, and C) morphs of Experiment 1.

## Bibliography

1. FW Campbell and JG Robson. Application of fourier analysis to the visibility of gratings. J. Physiol., 197(3):551–566, August 1968. ISSN 1469-7793,0022-3751. doi: 10.1113/jphysiol.1968.sp008574.

2. MS Banks, WS Geisler, and PJ Bennett. The physical limits of grating visibility. Vision Res., 27(11):1915–1924, January 1987. ISSN 0042-6989,1878-5646. doi: 10.1016/0042-6989(87)90057-5.

3. J Nachmias and A Weber. Discrimination of simple and complex gratings. Vision Res., 15(2):217–223, February 1975. ISSN 0042-6989,1878-5646. doi: 10.1016/0042-6989(75)90210-2.

4. D Regan and KI Beverley. Postadaptation orientation discrimination. J. Opt. Soc. Am. A, 2 (2):147–155, February 1985. ISSN 0740-3232,2375-1169. doi: 10.1364/josaa.2.000147.

5. MA Webster, KK De Valois, and E Switkes. Orientation and spatial-frequency discrimination for luminance and chromatic gratings. J. Opt. Soc. Am. A, 7(6):1034–1049, June 1990. ISSN 0740-3232,2375-1169. doi: 10.1364/josaa.7.001034.

6. DC Burr and SA Wijesundra. Orientation discrimination depends on spatial frequency. Vision Res., 31(7-8):1449–1452, 1991. ISSN 0042-6989,1878-5646. doi: 10.1016/0042-6989(91)90064-c.

7. Kathy T Mullen and William H A Beaudot. Comparison of color and luminance vision on a global shape discrimination task. Vision Res., 42(5):565–575, March 2002. ISSN 0042-6989,1878-5646. doi: 10.1016/s0042-6989(01)00305-4.

8. David J Tolhurst, Caterina Ripamonti, C Alejandro Párraga, P George Lovell, and Tom Troscianko. A multiresolution color model for visual difference prediction. In Proceedings of the 2nd symposium on Applied perception in graphics and visualization, pages 135–138, New York, NY, USA, August 2005. ACM. ISBN 9781595931399. doi: 10.1145/1080402.1080427.

9. Bruce C Hansen and Edward A Essock. A horizontal bias in human visual processing of orientation and its correspondence to the structural components of natural scenes. J. Vis., 4(12):1044–1060, December 2004. ISSN 1534-7362. doi: 10.1167/4.12.5.

10. Bruce C Hansen, Edward A Essock, Yufeng Zheng, and J Kevin DeFord. Perceptual anisotropies in visual processing and their relation to natural image statistics. Network, 14 (3):501–526, August 2003. ISSN 0954-898X,1361-6536. doi: 10.1088/0954-898x/14/3/307.

11. Yury Petrov. Luminance correlations define human sensitivity to contrast resolution in natural images. J. Opt. Soc. Am. A Opt. Image Sci. Vis., 22(4):587–592, April 2005. ISSN 1084-7529,1520-8532. doi: 10.1364/josaa.22.000587.

12. Uri Hasson, Rafael Malach, and David J Heeger. Reliability of cortical activity during natural stimulation. Trends Cogn. Sci., 14(1):40–48, January 2010. ISSN 1364-6613,1879-307X. doi: 10.1016/j.tics.2009.10.011.

13. Robert L Goldstone and Andrew T Hendrickson. Categorical perception: Categorical perception. Wiley Interdiscip. Rev. Cogn. Sci., 1(1):69–78, January 2010. ISSN 1939-5078,1939-5086. doi: 10.1002/wcs.26.

14. Nathan Destler, Manish Singh, and Jacob Feldman. Shape discrimination along morphspaces. Vision Res., 158:189–199, May 2019. ISSN 0042-6989,1878-5646. doi: 10.1016/j.visres.2019.03.002.

15. Masataka Sawayama, Yoshinori Dobashi, Makoto Okabe, Kenchi Hosokawa, Takuya Koumura, Toni P Saarela, Maria Olkkonen, and Shin’ya Nishida. Visual discrimination of optical material properties: A large-scale study. J. Vis., 22(2):17, February 2022. ISSN 1534-7362. doi: 10.1167/jov.22.2.17.

16. Katharine A Shapcott, Marvin Weigand, Mina Glukhova, Martha N Havenith, and Marieke L Schölvinck. DomeVR: Immersive virtual reality for primates and rodents. PLoS One, 20(1):e0308848, January 2025. ISSN 1932-6203. doi: 10.1371/journal.pone.0308848.

17. Keiji Matsuda, Takeshi Nagami, Yasuko Sugase, Aya Takemura, and Kenji Kawano. A widely applicable real-time mono/binocular eye tracking system using a high frame-rate digital camera. In Human-Computer Interaction. User Interface Design, Development and Multimodality, Lecture Notes in Computer Science, pages 593–608. Springer International Publishing, Cham, May 2017. ISBN 9783319580708,9783319580715. doi: 10.1007/978-3-319-58071-5_45.

18. Cynoclub. Emperor angelfish (pomacanthus imperator) stock image (image ID - 18739301). https://www.dreamstime.com/stock-image-emperor-angelfish-image18739301. Accessed: 2026-1-7.

19. Nawaj Panichphol. Discus fish stock photo (image ID 52328078). https://www.dreamstime.com/stock-photo-discus-fish-snaks-skin-thailand-image52328078. Accessed: 2026-1-7.

20. James Scott McDonald and Yoav Tadmor. The perceived contrast of texture patches embedded in natural images. Vision Res., 46(19):3098–3104, October 2006. ISSN 0042-6989,1878-5646. doi: 10.1016/j.visres.2006.04.014.

21. S Appelle. Perception and discrimination as a function of stimulus orientation: the “oblique effect” in man and animals. Psychol. Bull., 78(4):266–278, October 1972. ISSN 0033-2909,1939-1455. doi: 10.1037/h0033117.

22. KT Mullen, WH Beaudot, and WH McIlhagga. Contour integration in color vision: a common process for the blue-yellow, red-green and luminance mechanisms? Vision Res., 40(6):639–655, 2000. ISSN 0042-6989,1878-5646. doi: 10.1016/s0042-6989(99)00204-7.

23. Judy Borowski, Roland S Zimmermann, Judith Schepers, Robert Geirhos, Thomas S A Wallis, Matthias Bethge, and Wieland Brendel. Exemplary natural images explain CNN activations better than state-of-the-art feature visualization. arXiv [cs.CV], October 2020. doi: 10.48550/arXiv.2010.12606.

24. David G Jones, Nicole D Anderson, and Kathryn M Murphy. Orientation discrimination in visual noise using global and local stimuli. Vision Res., 43(11):1223–1233, May 2003. ISSN 0042-6989,1878-5646. doi: 10.1016/s0042-6989(03)00095-6.

25. Holly E Gerhard, Felix A Wichmann, and Matthias Bethge. How sensitive is the human visual system to the local statistics of natural images? PLoS Comput. Biol., 9(1):e1002873, January 2013. ISSN 1553-7358,1553-734X. doi: 10.1371/journal.pcbi.1002873.

26. MJ Morgan, AJ Mason, and S Baldassi. Are there separate first-order and second-order mechanisms for orientation discrimination? Vision Res., 40(13):1751–1763, 2000. ISSN 0042-6989,1878-5646. doi: 10.1016/s0042-6989(00)00015-8.

27. Harriet A Allen, Robert F Hess, Behzad Mansouri, and Steven C Dakin. Integration of first- and second-order orientation. J. Opt. Soc. Am. A Opt. Image Sci. Vis., 20(6):974–986, June 2003. ISSN 1520-8532,1084-7529. doi: 10.1364/josaa.20.000974.

28. Antonio Torralba and Aude Oliva. Statistics of natural image categories. Network, 14(3): 391–412, August 2003. ISSN 0954-898X,1361-6536. doi: 10.1088/0954-898x/14/3/302.

